# High photon count rates improve the quality of super-resolution fluorescence fluctuation spectroscopy

**DOI:** 10.1101/783639

**Authors:** Falk Schneider, Pablo Hernandez-Varas, B. Christoffer Lagerholm, Dilip Shrestha, Erdinc Sezgin, M. Julia Roberti, Giulia Ossato, Frank Hecht, Christian Eggeling, Iztok Urbančič

## Abstract

Probing the diffusion of molecules has become a routine measurement across the life sciences, chemistry and physics. It provides valuable insights into reaction dynamics, oligomerisation, molecular (re-)organisation or cellular heterogeneities. Fluorescence correlation spectroscopy (FCS) is one of the widely applied techniques to determine diffusion dynamics in two and three dimensions. This technique relies on the temporal autocorrelation of intensity fluctuations but recording these fluctuations has thus far been limited by the detection electronics, which could not efficiently and accurately time-tag photons at high count rates. This has until now restricted the range of measurable dye concentrations, as well as the data quality of the FCS recordings, especially in combination with super-resolution stimulated emission depletion (STED) nanoscopy.

Here, we investigate the applicability and reliability of (STED-)FCS at high photon count rates (average intensities of up to 40 MHz) using novel detection equipment, namely hybrid detectors and real-time gigahertz sampling of the photon streams implemented on a commercial microscope. By measuring the diffusion of fluorophores in solution and cytoplasm of live cells, as well as in model and cellular membranes, we show that accurate diffusion and concentration measurements are possible in these previously inaccessible high photon count regimes. Specifically, it offers much greater flexibility of experiments with biological samples with highly variable intensity, e.g. due to a wide range of expression levels of fluorescent proteins. In this context, we highlight the independence of diffusion properties of cytosolic GFP in a concentration range of approx. 0.01–1 μM. We further show that higher photon count rates also allow for much shorter acquisition times, and improved data quality. Finally, this approach also pronouncedly increases the robustness of challenging live cell STED-FCS measurements of nanoscale diffusion dynamics, which we testify by confirming a free diffusion pattern for a fluorescent lipid analogue on the apical membrane of adherent cells.

## Introduction

Fluorescence correlation spectroscopy (FCS) has, since its introduction almost 50 years ago, become a widely applied technique to study diffusion dynamics in synthetic and biological applications^1–3^. It has greatly contributed to the understanding of molecular diffusion in model systems and living cells, both in 2D (in vitro models or cellular membranes) and in 3D (solution or cellular cytoplasm and nucleus) environments^4–8^. Notably, it has offered fundamental insights into the dynamic organisation of living systems at the molecular level, e.g. by characterising the transient, dynamic, yet structured nature of the organisation of fluid membranes^4,5,9^.

FCS provides a plethora of information about molecular dynamics. The diffusion rates and local concentrations of fluorescent molecules can be determined directly from the autocorrelation functions^1,10,11^. The spatial variability can be further evaluated by laser-scanning^12–14^ and imaging-based variants of FCS^15–18^. Further, molecular interactions can be probed either directly, e.g. binding of molecules detected by cross-correlation (FCCS)^19^, or indirectly via variations of the apparent diffusion coefficient at different length scales, measured by spot-variation FCS^20^ providing information on diffusion modes as in single particle tracking^21^. Finally, the combination of FCS with super-resolution stimulated emission depletion (STED) microscopy allows direct observation of nanoscale diffusion dynamics, shedding new light on molecular organisation below the diffraction limit^22^.

All these invaluable details are extracted from intensity fluctuations due to the transit of fluorescent molecules through the observation spot of the microscope. As the fluctuations (i.e. bursts in the fluorescence intensity trace) are most obvious for sparsely labelled samples, FCS is often considered a single molecule technique, and has thus been shown multiple times to perform accurately in the range of pico- and nanomolar concentrations^2^. These concentrations, though, can be far from physiological levels present in living systems, where molecular abundance can be much higher (e.g. average concentration of a protein in eukaryotic yeast cells is estimated to be around 1 µM^23^). Nevertheless, it has been theoretically predicted and experimentally verified that FCS can perform similarly and can generate accurate results also for much larger concentrations (>100 nM)^24^. In this regime, the main factors for signal quality of FCS, often described by the signal-to-noise ratio (SNR), are the acquisition time (*T*), and the number of detected photons per molecule (i.e. molecular brightness, *B*, which depends on the absorption cross section and quantum yield of the dye, the power of the excitation laser, and the detection efficiency of the measurement setup): SNR ∝*B* × *T*^1/2^ (see for example^11,25–28^).

For the most efficient and reliable detection of fluorescence fluctuations, sensitive single-photon-counting detectors are typically used, often coupled to fast electronics that enable accurate recording of photon arrival times thus also allowing additional photon filtering in post-processing^29^. One of the main drawbacks of this instrumentation, however, has been its rather long dead time after each photon detection (>100 ns)^30^, limiting photon count rates to a few MHz, which is far lower than the typical repetition rate of excitation lasers running at 20–80 MHz. This has posed a severe limitation to the accuracy and flexibility in FCS experiments at high fluorophore concentrations, which are however unavoidable for many applications – for example when measuring binding dynamics of low affinity, or diffusion dynamics and concentrations of cellular proteins at different expression levels. Several approaches have been developed to enable FCS measurements even in such cases: labelling of only a fraction of the molecules, reduction of the simultaneously visible fluorophores via fluorescence photoswitching^31,32^, splitting-up of the signal onto several detectors such as on custom-built detector banks^33^, or reduction of the effective observation volume^34,35^ using for example small sample containers^36^, near-field structures^37,38^, plasmonic near-field optics^39–41^, or super-resolution STED microscopy^5,42^. Unfortunately, all of these techniques introduce more complexity and possible bias, for example due to required controls to check whether the fraction of labelled or photoswitched molecules truly reflects the entire population, influence on the sample and fluorescent molecules by surface or small volume effects, setup complexity, or perplexing photophysics of the fluorescent label.

Here, we demonstrate the straightforward realisation of FCS measurements at high photon count rates on a commercially available microscope, using novel photon counting instrumentation. By measuring the diffusion of fluorescent dyes in solutions, artificial and cell membranes, we explore performance, capabilities, accuracy, applicability and limitations of confocal FCS and STED-FCS experiments at photon count rates of up to 20–30 MHz per detection channel, revealing great potential for FCS experiments at high count rates. As examples, we show the independence of cytosolic protein diffusion on cellular expression levels and the application of high count rates to STED-FCS measuring the diffusion behaviour of lipids in apical cellular plasma membranes.

## Results and Discussion

### Non-saturated photon detection at high dye concentrations and laser excitation powers

We first tested the advanced photon counting instrumentation, implemented on a confocal and STED-capable microscope, by recording fluorescence fluctuation data from a single dye (Atto655, chosen for its low population of triplet states) diffusing in aqueous solution at different concentrations or excitation laser powers, resulting in different photon count rates. The photon counting instrumentation included hybrid detectors with very short dead times and fast FPGA electronics with real-time GHz sampling and pattern matching, which together with the 80 MHz pulsed fluorescence excitation allows for detection of photon count rates of tens of MHz without corrections (as described in detail in the Materials and Methods section). Figure 1A and B show fluctuations in the normalized photon count rates over time as recorded for three different dye concentrations and laser excitation powers, respectively. As expected from theory^10,25–28^, the relative fluctuations around the average count rate decrease with increasing dye concentration, but much less so with excitation laser power. Most importantly, we could follow a linear increase of photon count rate with dye concentration and laser excitation power (Figure 1C and D), as expected in the absence of limitations in detection electronics. Note that approximate linearity was maintained despite employing dye concentrations of up to 1 µM and registering photon count rates of up to 20-30 MHz. The non-linearity introduced at excitation laser powers larger than 40 µW (≈ 30 kW/cm^2^, Figure 1D) were to be expected due to dye photobleaching and saturation of excited state population and consequently fluorescence emission (i.e. not due to detector saturation, compare Figure 1C at same count rate levels), while slight saturation effects at very high dye concentrations may result from photon re-absorption and dye self-quenching, as indicated previously^33,43^. In due consideration of acquisition count rate being the limiting factor in conventional equipment, we present the rest of the data as a function of this parameter.

**Figure 1:**
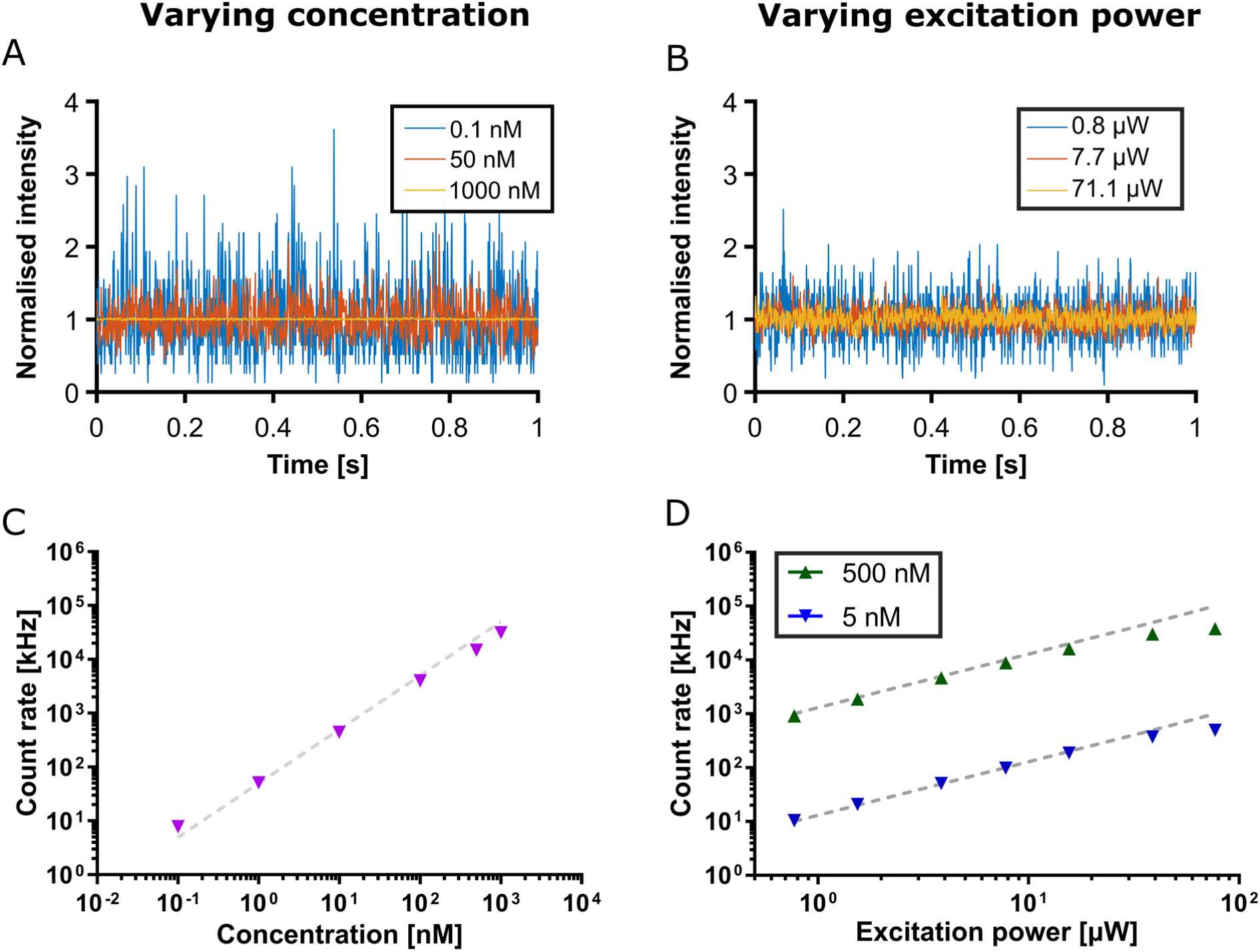
Influence of varying concentration and excitation laser power on FCS measurements of the dye Atto655 in aqueous solution. **A**,**B** Normalized detected photon count rate data (intensity) over time at different dye concentrations (A, 0.1 nM, 50 nM, and 1 µM as labelled; excitation laser power 15.4 µW) and excitation laser powers (B, as labelled; concentration 5 nM), highlighting the reduction in relative intensity fluctuations for higher concentrations and excitation laser powers. **C**,**D** Detected photon count rates versus instituted dye concentration (C, dashed lines indicate the expected linear increase; excitation laser power 15.4 µW) and instituted excitation laser power for 5 nM (blue) and 500 nM (green) (D, dashed lines indicate the expected linear increase). Values are averages of three repetitions (acquisition time 15 s each), and standard deviations are smaller than the size of the symbols.

### FCS noise levels at different count rates

The ability to record photon time traces at high count rates consequently allowed us to acquire FCS data for Atto655 up to 1 µM high dye concentrations and excitation laser powers up to 40 µW (≈ 30 kW/cm^2^). The autocorrelation curves for these unconventional conditions show similar decays as for low dye concentrations and laser powers (Figure 2A and B). From common FCS theory (assuming large particle concentrations and excluding non-linear photo-physical effects such as saturation and photobleaching), FCS-derived average transit times should be independent of dye concentration and laser power, while the amplitude of the autocorrelation curve should linearly decrease with dye concentration and stay constant with laser power. We could well recover this behaviour from fitting our FCS data, even when recorded at count rates of up to 20-30 MHz (Figure 2C and D). As before, deviations at the highest tested laser powers >40 µW (primarily apparent as a drop in the values of the transit time and FCS amplitudes, Figure 2D), can be attributed to dye photobleaching and fluorescence emission saturation. For dyes other than Atto655, excessive pumping into their triplet states would result in additional deterioration at high excitation powers and would thus need to be carefully considered as well^44,45^.

**Figure 2:**
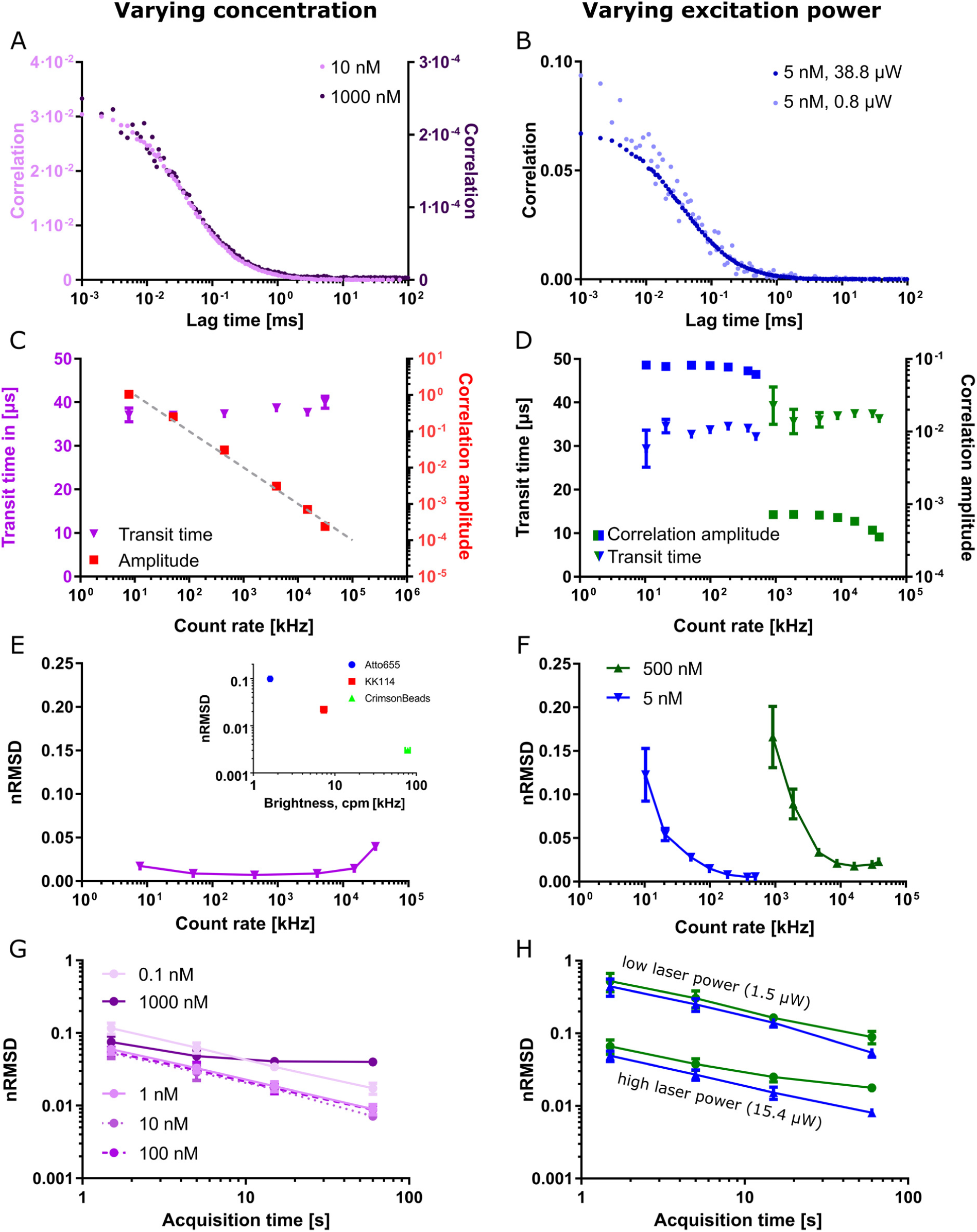
Detailed analysis of the FCS data of Atto655 in aqueous solution at varying instituted dye concentration c_dye_ (left panels) and excitation laser power P_laser_ (right panels). **A**,**B** Representative FCS curves at different concentrations (A, as labelled, P_laser_ = 15.4 µW) and excitation laser powers (B, as labelled, c_dye_ = 5 nM). **C-F** Transit time, correlation amplitude (C and D, axes and colours as labelled, D: for two c_dye_ as labelled), and nRMSD (i.e. noise in correlation data, E and F; F: for two c_dye_ as labelled) as determined from FCS for different photon count rates, i.e. different c_dye_ (C, E) and P_laser_ (D, F). (**Inset E**) Values of nRMSD as determined from FCS for different _dye_s in aqueous solution with different brightness, i.e. counts-per-molecule (cpm) (average over c_dye_ ≈ 0.01–1 µM or 1:100–1:2000 dilutions of the stock bead suspension (see Figure SI 3), P_laser_ = 1.5 µW). **G**,**H** Values of nRMSD as determined from FCS data of Atto655 at different acquisition times and at varying c_dye_ (G, P_laser_ = 15.4 µW) and P_laser_ (H, c_dye_ = 5 or 500 nM) as labelled. Data points represent averages and standard deviations (unless smaller than the symbols) of three repetitions (60 s acquisition time if not indicated otherwise).

An interesting feature of the FCS data recorded at different dye concentrations or excitation laser powers, i.e. photon count rates, are the different noise levels and resulting data quality. From theory^11,24,26^, the noise in FCS data should linearly decrease with increasing excitation laser power and be independent of dye concentration. The latter has for example been experimentally verified for dye concentrations of up to around 100 nM^24^. Consequently, we set out to investigate noise levels for FCS data recorded at the large dynamic range of dye concentrations and laser powers, which became accessible using the new equipment.

Already visual inspection of our FCS data recorded at the different conditions (Figure 2A and B) indicated notable differences in noise levels (shown as the spread of the correlation curve), especially for the different excitation laser powers, which was most pronounced at short lag times (in this case up to around 40 μs, roughly corresponding to the transit time of the dye). The non-trivial estimation of noise levels in FCS data has been the subject of intense investigations^11,24–29^. In our experience, the noise levels in the current FCS data were well estimated by the root-mean-square of the fitting residuals normalized to the amplitude (nRMSD, calculated for short lag times up to the transit time), with lower values indicating better data quality (note that nRMSD was not used as the residuals minimisation metric in our fitting protocol; see Materials and Methods and Figure SI 1 for details). The nRMSD provided us with a single value for the data quality of each measurement. Conveniently, this measure also roughly corresponded to the relative standard deviation of the values of the average transit time as determined from fitting of the data (Figure SI 2, the nRMSD relates closely to the measurement error, i.e. the experimental standard deviation of the correlation at every data point, Figure SI 1). In accordance with the theoretical predictions^11,24,26^, the noise in the FCS data and thus nRMSD were only weakly affected by varying concentration (Figure 2E), but could be greatly improved by increased excitation laser power (Figure 2F). The excitation laser power directly increases the dyes’ excited state population and thus fluorescence emission rate and the average detected count rate per single dye (molecular brightness), which is the reason for the improvements in noise levels. Under comparable measurement conditions, the nRMSD is therefore also a direct indicator of the molecular brightness of the investigated dye (inset in Figure 2E and Figure SI 3). Deteriorated noise levels, i.e. higher nRMSD values, were again observed at dye concentrations around 1 µM, but were much less pronounced at the highest excitation laser powers above 20–40 µW, despite saturation of photon count rates due to photophysical limitations of the dye (Figure 1D) and deviations of values of the transit times and correlation amplitudes (Figure 2D) as highlighted above.

### FCS noise levels at different acquisition times

Predicted from theory and to a certain extent verified experimentally^11,24,26^, the noise in FCS data should decrease with the square root of the acquisition time. We could well reproduce this dependence for different dye concentrations and excitation laser powers (Figure 2G and H; again, the same issues as outlined above caused deviations at high dye concentrations and excitation laser powers). This data establishes unique possibilities of adapting to experimental conditions. Due to the absence of saturation effects in photon count rates in the 5–500 nM concentration range, the possibility of increasing the laser power and detecting correspondingly higher photon count rates does not only increase the data quality (i.e. lower nRMSD values), but alternatively allows for a significant reduction (up to two orders of magnitude) in the acquisition time required for generating similar data quality (Figure 2G and H, Figure SI 2).

### FCS of cytosolic GFP in live cells at various expression levels

The possibility of acquiring accurate FCS data in a wide range of dye concentrations allows simplification or realization of experiments under challenging conditions. For example, FCS-based measurements of diffusion or concentration of fluorescent proteins in cells are usually challenged by the naturally varying expression levels of the fluorescent proteins, as shown in the representative confocal image in Figure 3A for HeLa cells expressing cytoplasmic GFP (green fluorescent protein). Using standard FCS instrumentation, only carefully chosen dim cells would be measurable, which may represent only a small, and not necessarily representative fraction of all cells, introducing a potential source of bias.

**Figure 3:**
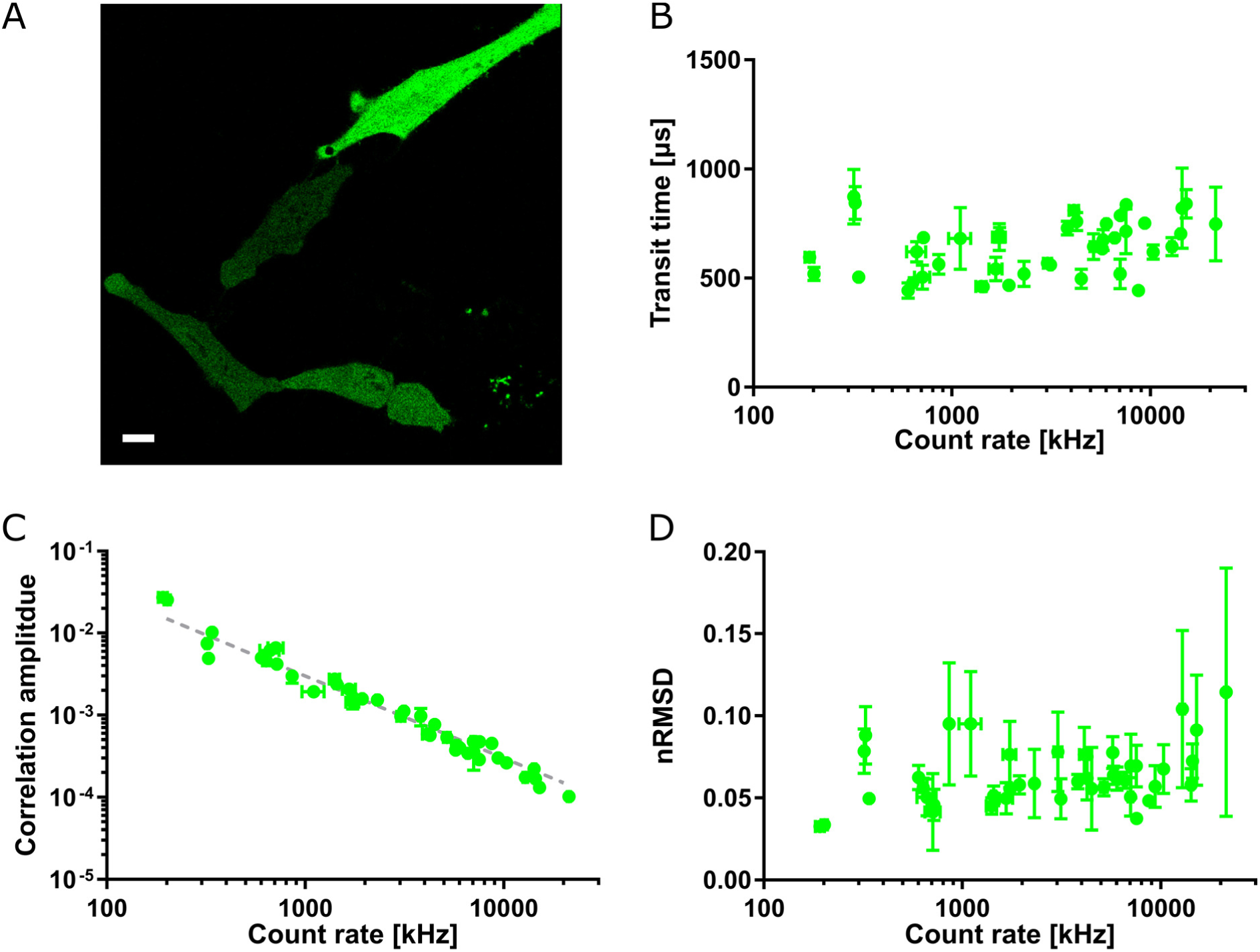
Acquiring FCS data in live cells at different expression levels of the cytosolic fluorescent protein GFP-SNAP. **A** Representative confocal image of HeLa cells transiently transfected with cytoplasmic GFP-SNAP, highlighting different brightness and thus expression levels. Scale bar: 10 µm. **B-D** Values of transit time (B), correlation amplitude (C, with the grey guideline indicating the predicted inverse relationship), and nRMSD (i.e. noise in correlation data, D) as determined from FCS for different photon count rates, originating from different expression levels. Averages and standard deviations of at least three measurements per cell (acquisition time 15 s, excitation laser power 0.6 µW).

Using our current setup, we could now record FCS data for all HeLa cells irrespective of their fluorescence intensity, revealing average transit times of GFP over a wide range of photon count rates and thus concentrations resulting from different expression levels (Figure 3B). The photon count rates are correlated with the correlation amplitude (Figure 3C), which is inversely proportional to the average number of fluorescent molecules in the observation volume (see Materials and Methods) and thus concentration and expression level of GFP can be inferred. Taking our observation volume of about 1 fL, we can estimate concentrations of GFP of approx. 0.01–5 μM between the differently expressing cells (see Materials and Methods). These data indicate that within the tested range the mobility of GFP is independent of expression level. In addition, the quality of the FCS data as quantified by the nRMSD values was maintained over the range of tested expression levels (Figure 3D), as predicted from the behaviour of the organic dye in solution (compare Figure 2). Only in the regime beyond 10 MHz (in our case corresponding to concentrations around 1 µM), we observed a slight signal deterioration due to, for example, possible out-of-focus and self-absorption contributions, reflected in an increase of the nRMSD (Figure 3D) and a decrease in the amplitude beyond the predicted inverse relationship with the count rate (grey dashed guide line in Figure 3C).

### STED-FCS of lipid dyes in model membranes

The main strength of FCS on a super-resolution STED microscope, STED-FCS, is the ability to directly report on nanoscale molecular mobility and thus determine apparent values of diffusion coefficients from the average transit times (see Materials and Methods) for different observation spot sizes – from conventional confocal spot sizes with lateral diameters of around 200 nm, down to STED microscopy recordings with observation spot diameters of 30–40 nm. From the dependency of the apparent values of the diffusion coefficient on the observation spot diameter, STED-FCS has provided insights into the molecular diffusion modes, similar to spot-variation FCS^20^, but now at the relevant molecular scale, which is particularly valuable for the elucidation of the nanoscale architecture of biological membranes^22^. However, measurements at various sizes of the effective observation spot inherently impose a large variation in the average number of fluorescent molecules in the observation spot (*N*) and thus detected photon count levels (Figure 4A, schematics). Large observation spots at the confocal recordings entail already high count rates and high values of *N* at rather low dye concentrations, while the smaller observation spots at the STED microscopy recordings require relatively large dye concentrations to reach photon count rates and values of *N* that are high enough for allowing reasonably low acquisition times (it has also been shown theoretically that too low count rates or concentrations lead to noisy and inaccurate FCS data^11,24,26^). This has limited the range of useful dye concentrations in STED-FCS measurements using conventional detection electronics.

**Figure 4:**
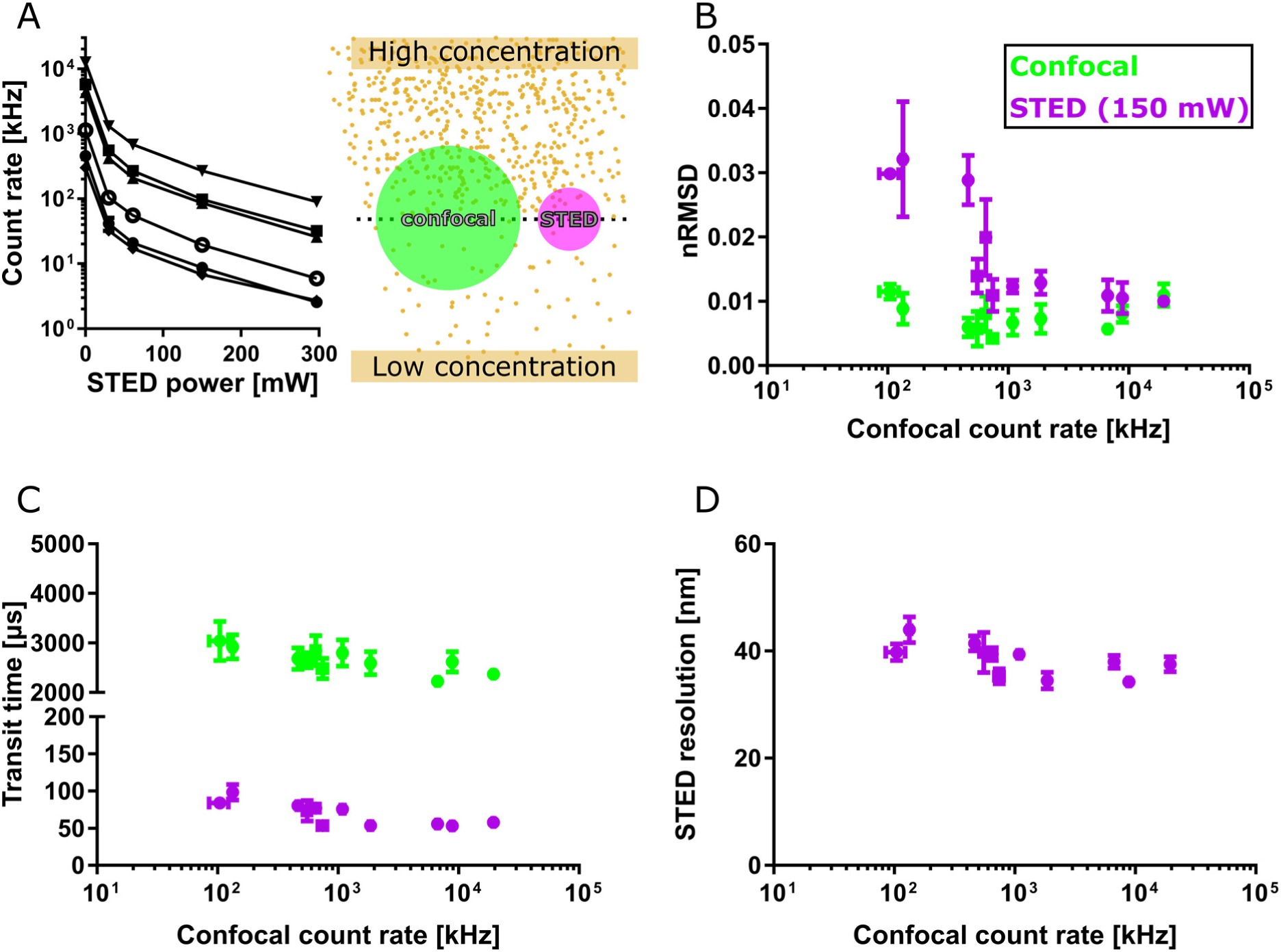
STED-FCS in supported lipid bilayers (SLBs) at various dye concentrations. **A right** Sketch illustrating the shrinking of the observation spot from the confocal (green) to the STED (purple) modus for a high or low concentration of dye molecules (yellow dots). **Left** Average fluorescence intensity experimentally recorded for different concentrations of a fluorescent lipid analogue (Abberior STAR Red-DPPE; different symbols) in DOPC SLBs for increasing powers of the STED laser. From these data, the noise level of FCS data (nRMSD, **B**), the average fitted transit time (**C**), and calculated apparent STED resolution (**D**) for confocal (green) and STED-FCS recordings (purple, 150 mW depletion power) are plotted against the respective confocal acquisition count rates, which are the limiting factor in each STED-FCS experiment. Excitation power 2.3 µW, acquisition time 15 s. Each data-point represents the average and standard deviation of at least three measurements per SLB preparation.

To evaluate the performance of the new detection electronics in STED-FCS, we recorded data at varying concentrations of the fluorescent lipid analogue Abberior STAR Red-DPPE (DPPE: 1,2-dipalmitoyl-sn-glycero-3-phosphoethanolamine) diffusing in a fluid supported lipid membrane bilayer (SLB, composed of lipids DOPC, 1,2-dioleoyl-sn-glycero-3-phosphocholine), which is a convenient and well characterised model membrane system, often used as a control sample in STED-FCS experiments^22^. While increasing the lipid analogue concentration and pushing the confocal count rates beyond the conventional FCS range (Figure 4A) did not influence the noise in confocal measurements (Figure 4B, green data points), it - as expected - significantly improved the data quality of the STED-FCS recordings (low nRMSD values, Figure 4B, purple data points; note that for plotting of these, the respective confocal count rates, which limit the experimental conditions for STED recordings, were used as the x-value). Also as expected from theory, the values of transit times were smaller in the STED compared to the confocal recordings (due to the reduced observation spot size in the STED mode, here roughly 40 nm in diameter) and hardly changed with instituted concentrations of the fluorescent probe (i.e., at increased count rate, Figure 4C). Similarly, the observation spot diameters as determined from the recorded FCS data remained constant (Figure 4D, see Materials and Methods for details about its calculation), all in all highlighting the great flexibility and improvement in STED-FCS experiments when employing non-saturating detection electronics.

For simplicity, we demonstrated the above effects at a single STED laser power (150 mW, observation spot diameter of 40 nm), but the conclusions held also true for other STED laser powers (and thus observation spot sizes). Similar or even larger improvements of STED-FCS data quality as by increasing the dye concentration were achieved by increasing the excitation laser power (Figure SI 4; note that we, in contrast to the previous data of Atto655, now included an additional decay due to triplet state population in FCS data model, see Materials and Methods and Figure SI 4E). Effects due to saturation of the excited states (such as the triplet state) of the fluorescent label started to deteriorate the signal quality and bias the extracted parameter values only at laser powers beyond 25 μW, supporting the benefits of high count rates for STED-FCS experiments with the excellent dyes available nowadays.

### STED-FCS in live cell membranes

Finally, we verified the reliability of STED-FCS measurements at high photon count rates for measurements in living cells. We labelled the plasma membrane of live HeLa cells using the fluorescent lipid analogue Cholesterol-PEG-Abberior STAR Red (Chol-PEG-KK114, Figure 5A), for which previous studies have consistently indicated free diffusion in cellular plasma membranes^46^. The resulting STED-FCS data show low nRMSD values, i.e. low noise levels in the correlation data, which can in the STED microscopy mode be significantly improved to almost confocal quality by increasing either the excitation laser power or concentration of Chol-PEG-KK114 (i.e. total count rate, Figure 5B) without biasing the resulting values of transit times (Figure 5C).

**Figure 5:**
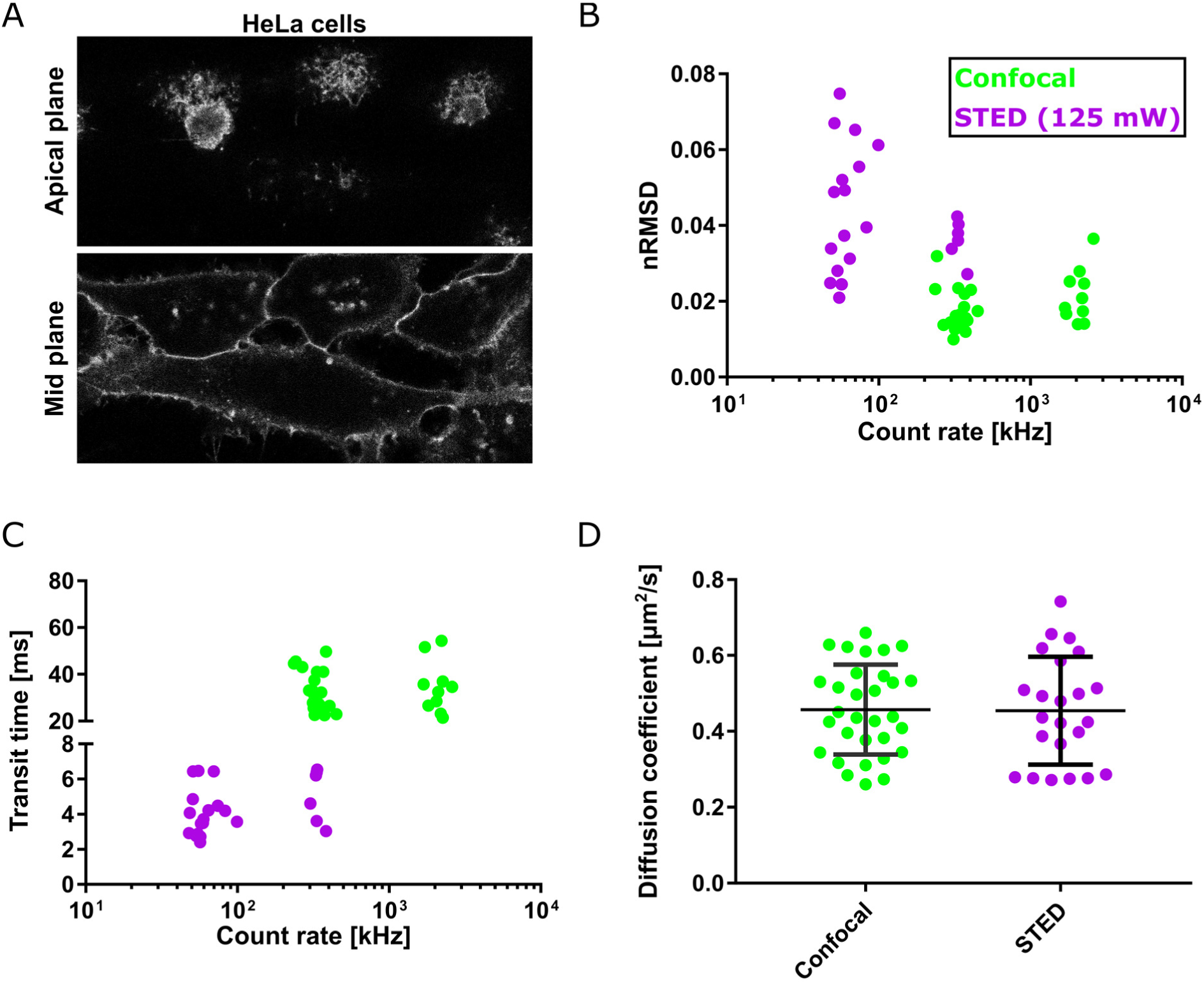
STED-FCS of Abberior STAR-Red-PEG-Cholesterol in membranes of live HeLa cells. **A** Confocal image of the apical plane (roughly 10 µm above the cover slip) and mid plane of HeLa cells, membrane labelled with the fluorescent cholesterol analogue (image sizes 60 × 27 µm^2^) **B** The noise level of FCS data (nRMSD values) and **C** average fitted transit time for confocal (green) and STED-FCS recordings (125 mW depletion laser power, purple), in the top membrane of HeLa cells, plotted against their respective count rates. **D** Diffusion coefficient determined from FCS data in confocal and STED mode in the top membrane of the HeLa cells. Every dot represents a single FCS measurement. Excitation laser 2.3 µW, acquisition time 15 s.

Note that we here measured the diffusion in the apical membrane of HeLa cells, i.e. 5–10 μm above the microscope coverslip (Figure 5A), rather than in the basal membrane as before^46^. This avoids potential biasing effects by the coverslip surface. Yet, penetration through the aqueous cellular environment over such a distance causes spherical aberrations when employing a traditional oil-immersion STED microscope objective (due to the refractive index mismatch between water and oil)^47^, having detrimental effects on STED-FCS experiments (Figure SI 5). Such aberrations can either be corrected for using adaptive optics^48,49^ or employing a water immersion objective^50^, which shows constant signal levels and performance of for example STED and STED-FCS experiments in a wide range of focal depths (5–100 µm above the coverslip) without the need for depth-dependent readjustments of the correction collar (Figure SI 5). Taking the observation spot size as determined from calibration data (Figure SI 6, diameter of 100 nm), we could calculate apparent values of the diffusion coefficient (*D*) for both the confocal and STED mode (spot diameters of 280 and 100 nm, respectively; see Materials and Methods), which were both in the same range (*D* ≈ 0.45 µm^2^/s, Figure 5D) as observed before for the basal membrane^46^, highlighting free diffusion and indicating similar diffusion characteristics of the probe in the apical and basal plasma membrane^46^, i.e. negligible bias by the coverslip surface.

## Conclusions

We systematically evaluated the reduction in error and bias of FCS measurements recorded at high photon count rates, as enabled by novel detection electronics integrated into a turn-key microscope. We were able to record highly accurate FCS data with detected photon count rates of up to about 10 MHz, i.e. dye concentrations up to 1 μM. This improved performance introduces huge flexibility for performing FCS experiments to measure diffusion or concentration, previously impossible due to limitations in the detection electronics (allowing e.g. only recordings of photon count rates of up to few MHz). This now enables: 1) FCS measurements at high dye concentrations for e.g. low-affinity binding assays, 2) the recording of fluctuation data with reduced acquisition times by increasing the excitation power to higher count rates (dye photophysics permitting), 3) performing live-cell experiments in a wide range of expression levels of fluorescently tagged proteins, and 4) optimization of the data quality of STED-FCS recordings over a wide range of observation spot sizes by increasing dye concentration and/or excitation laser power. Using these features we could for example show that cytosolic diffusion of GFP was independent of expression level in live HeLa cells, and that the fluorescent lipid analogue was diffusing freely in the apical membrane similarly as reported before for the basal membrane^46^.

Improved detection instrumentation as the one presented here are becoming increasingly available and will be further optimized, pushing the ease of use of FCS or related measurements, such as fluorescence cross correlation spectroscopy (FCCS)^51^, fluorescence lifetime correlation spectroscopy (FLCS)^52^, number and brightness (N&B) analysis^53^, or line- and raster-scanning correlation spectroscopy (RICS)^12,54^. In combination with high-throughput methods this could enable the systematic evaluation of overexpression of fluorescent proteins^55^, tracking of dynamically changing diffusion properties, or other previously unattainable applications.

## Acknowledgements

The authors thank for the funding from MRC Proximity to Discovery funds (MC_PC_16082 (P2D Technologies 2 Therapies)), Marie Skłodowska-Curie Actions (707348; I.U.), Newton-Katip Celebi Institutional Links grant (352333122, E.S.), Medical Research Council (MC_UU_12010/unit programs G0902418 and MC_UU_12025), MRC/BBSRC/EPSRC (MR/K01577X/1), Wolfson Foundation, Wellcome Trust (104924/14/Z/14), Deutsche Forschungsgemeinschaft (Research unit 1905, Excellence Cluster Balance of the Microverse, Collaborative Research Centre 1278 Polytarget), Wellcome Institutional Strategic Support Fund, and Oxford internal funds (EPA Cephalosporin Fund and John Fell Fund), support from the Micron Oxford Advanced Bioimaging Unit (Wellcome Trust funding 107457/Z/15/Z), and Dr. Katharina Reglinski for kindly providing the GFP-SNAP plasmids.

## Conflict of interest

The authors declare no conflict of interest but need to note that MJR, GO and FH are employed at Leica Microsystems that manufactures and sells the SP8 STED FALCON used throughout the study.

## Materials and Methods

### Preparation of dyes in solution

Atto655 NHS-ester (AttoTec), Abberior STAR Red NHS-ester also termed KK114 (Abberior), and 20-nm crimson beads (Thermofisher) were stored at concentration >10 µM and diluted in PBS for measurements.

### Preparation of Supported Lipid Bilayers (SLBs)

SLBs were prepared by spin coating as described previously^56^. Briefly, a solution of 1 mg/mL DOPC (1,2-dioleoyl-sn-glycero-3-phosphocholine, Avanti Polar Lipids) dissolved in 1:2 Methanol:Chloroform was spin coated at 3200 rpm for 45 seconds on a 25 mm diameter cover slip. The formed lipid film was rehydrated with SLB buffer (150 mM NaCl, 10 mM HEPES, pH 7.4) and washed several times. All cover slips for SLB preparation were piranha cleaned (3:1, H_2_S0_4_:H_2_O_2_) and stored in water. SLBs were labelled with varying amounts of Abberior STAR Red-DPPE (1,2-dipalmitoyl-sn-glycero-3-phosphoethanolamine, Abberior).

### Tissue Culture

HeLa cells were cultured at 37 °C at 5% CO_2_ in high-glucose DMEM (Thermofisher) supplemented with 10 % FBS (Thermofisher), L-Glutamine (Thermofisher) and penicillin/streptomycin (Thermofisher). Cells were seeded onto 35 mm IBIDI glass bottom dishes coated with fibronectin (10 µg/mL for 5 min and washed with PBS) 24 h prior to performing the measurements.

CHO K1 cells were grown at 37 °C at 5% CO_2_ in DMEM/F12 (Lonza) supplemented with 10 % FBS and L-Glutamine (both Sigma), and labelled in L15 with a fluorescent lipid analog at room temperature. The transfections of GFP-SNAP, cytoplasmic GFP, obtained from Dr. Katharina Reglinski, were performed with Turbofect (ThermoFisher) according to the manufacturer’s protocol.

### Preparation of Giant Plasma Membrane Vesicles (GPMVs)

GPMVs were prepared as described previously^56,57^. In brief, HeLa cells were cultured as described above but seeded on 35 mm plastic bottom petri dishes. At a confluency of about 75 %, the cells were washed with GPMV buffer (150 mM NaCl, 2 mM CaCl_2_, 10 mM HEPES, pH 7.4) and then incubated with 25 mM PFA and 10 mM DTT in GPMV buffer for 2 hours at 37 °C. The GPMV containing supernatant was collected and labelled with Abberior STAR Red-PEG(2kDa)-Cholesterol (Abberior) at a final concentration of 0.5 μg/mL for 10 minutes. GPMVs were non-specifically immobilised on poly-*L*-lysine (PLL) coated surfaces as described before^14^. All diffusion measurements in GPMVs were performed on the top membrane.

### Instrumentation & microscopy

All experiments were performed on a Leica SP8 STED FALCON (Leica Microsystems) equipped with the HC PL APO 100x/1.40 Oil STED WHITE oil immersion objective lens (SLB measurements) and the HC PL APO 86x/1.20 W motCORR STED WHITE water immersion objective lens with a motorised correction collar (for solution, cytosolic, and apical cell membrane measurements). The STED WHITE 86x water lens has a working distance of 300 µm, and the motorised correction collar adjusts for refraction index mismatch by optimizing the signal for every sample (coverslip). It is worth noting that a single initial setting of the correction collar was sufficient to correct for varying depth over the investigated range of 100 µm (Figure SI 5). We used the 488-nm and 633-nm lines of a white light laser as the excitation source. STED-FCS experiments were performed using a 775 nm pulsed laser (80 MHz) for depletion with laser powers between 0 and 300 mW measured at the objective. STED delay time was optimised using an SLB sample and minimising the detected photon count rate under high-power STED illumination. The respective notch filters (775 nm, 633 nm or 488 nm) were used for emission clean up. For all measurements with constant excitation power we stayed below saturation intensity (by checking proportionality of excitation laser power and fluorescence intensity) as triplet pumping may result in an additional source of deviations^58^. Fluctuations in laser intensities, which in some other studies reflected in a pronounced correlation component with decay times on the order of seconds and had to be corrected for^33^, were not observable in our case – all FCS curves converged to 0 (see examples in Figures 1, SI 1, and SI 7).

All FCS experiments were performed using the hybrid detectors (HyD-SMDs), featuring very short dead times, and FALCON electronics allowing acquisition of TCSPC data at photon count rates of up to 80 Mcps per detection channel without the necessity for corrections, becoming comparable to or exceeding the repetition rates of commonly used pulsed excitation lasers. This implementation is based on sampling the signal from the pulsed laser and detectors using fast FPGA electronics and applying pattern matching to the resulting bitstreams, producing as output the photon arrival times with a resolution of 97 ps and dead time <1.5 ns, at GHz sampling rates (for more technical details, please see the Leica Falcon Application Note^59^). Though certain other detector types such as avalanche photodiodes (APDs) offer 2–3-fold higher quantum efficiency in the far-red part of the spectrum compared to HyD-SMDs, their dead times are typically around 30 ns, i.e. 20-fold longer. The here-employed technology thus offers the highest currently achievable overall count rates, which are now on pair with the repetition rate of the excitation laser. Nevertheless, future developments towards increasing the detectors’ quantum efficiencies will allow further benefits, e.g. the use of lower excitation laser powers to achieve similarly high count rates or superior signal-to-noise ratios at high STED powers.

Measurement times ranged as indicated from 10 to 60 seconds. Only for the lifetime measurements we used 40 MHz pulsing of the white light laser for excitation.

### Data analysis

Correlation, time trace cropping, gating and fitting was performed using the built-in routines in LAS-X (Leica Microsystems). Time gates were applied in STED-FCS to remove confocal or laser scattering contributions and therefore improve resolution (see Figure SI 7), while not affecting confocal measurements (Figure SI 7; marginal deterioration of signal-to-noise or slightly larger spread of the fitted parameters were barely noticable). Solution and cytoplasmic GFP data were fitted with a free 3D diffusion model including offset and a triplet component^44^ as appropriate (triplet correlation decay time of GFP 40 µs with relative amplitude fixed to 14 %, Atto655 no triplet population^60^, Abberior STAR Red triplet correlation decay time 5 µs with triplet amplitudes within 5–10 %; crimson beads triplet correlation decay time 10–100 µs with triplet amplitudes 3–5 %). SLB, GPMV and cell membrane data were fitted with a 2D anomalous diffusion model^61^ (including offset and triplet time for Abberior STAR Red-PEG-Chol or -DPPE: 5 µs with relative amplitude around 10 %, and up to 30 % at the highest excitation powers). The parameter optimisation was performed using a Levenberg-Marquardt non-linear least-squares minimisation method with data-points weighted by their standard deviations, which were estimated from variations in ACFs calculated from sub-sections of the intensity time trace^27^. Some representative autocorrelation curves together with their fits, non-weighted fitting residuals, and reduced *χ*^2^ values^27^ are displayed in Figures SI1 and SI8.

As a measure of data quality and curve smoothness, defining the precision of the extracted fitted parameters, nRMSD values were calculated by taking the root-mean-square difference between the measured FCS curve and its fit up to the transit time, and normalised to the fitted amplitude. Such measure yielded similar values as the experimental standard deviation of the autocorrelation curves^27^ normalised to the amplitude (Figure SI1 and SI8; for easier comparison to nRMSD, the main data quality assessment tool throughout this work, we there display non-weighted residuals, but plot also standard deviations to put the residuals in the right perspective relevant to fitting). Smooth data-points at longer lag times were excluded, to avoid the influence of model miss-fitting in extreme conditions applied in this study, e.g. when curves were distorted due to saturation or photobleaching effects at the highest excitation powers, or by random bright transits occurring in cells. The constructed measure nRMSD thus correlates well with the relative standard deviation of the fitted transit times (Figure SI 2D).

For concentration estimation, we assumed a confocal volume of 1 fL. Given that the correlation amplitude relates to the inverse average number of particles in the focal volume, concentrations can be estimated^10^. For STED-FCS experiments the diffusion coefficient was calculated as described before^22^ using the following formula:

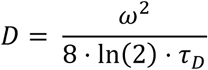

Where *D* is the apparent diffusion coefficient (in confocal or STED), *ω* refers to the full width half max (FWHM) of the observation spot (in confocal or STED) and *τ*_*D*_ to the transit time extracted from the FCS fit. The FWHM was determined by confocal and STED imaging of fluorescent beads (20 nm crimson beads). Using the assumption that the fluorescently labelled lipids diffuse freely in a SLB, the FWHM can be calculated as a function of STED power using the following equation^22^:

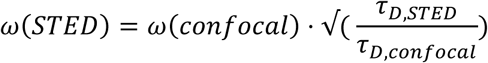

Note that we used the top membrane of immobilised GPMVs labelled with Abberior STAR Red-PEG-Cholesterol to determine the FWHM far away from the surface using a water immersion objective.

## Supporting figures

**Figure SI 1:**
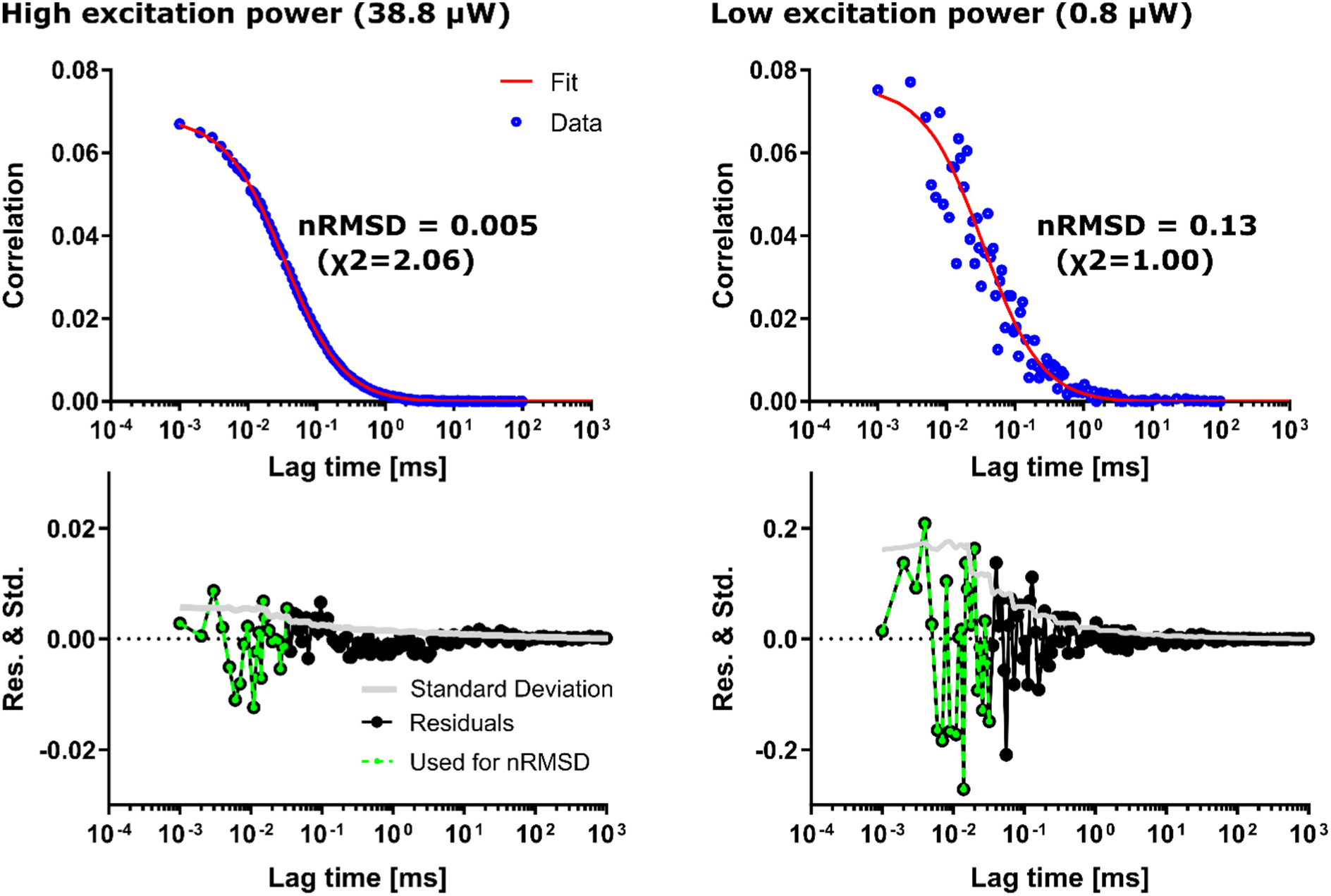
Calculation of nRMSD values from the (non-weighted) fitting residuals as a measure of the noise level in autocorrelation curves. FCS data of 5 nM Atto655 in aqueous solution acquired at high and low excitation power (38.8 and 0.8 µW, blue data-points in left and right panels, respectively) with respective fits (red curves) and fitting residuals (black) and experimental standard deviations of the autocorrelation at every lag time (grey), both normalised by the amplitude, below. The residuals highlighted in green (up to the lag time of the fitted transit time) where used for the calculation of the nRMSD value; the lower the nRMSD, the better the data quality. In fitting, though, the residuals were weighted by the respective standard deviations for the determination of the reduced χ^2^-values as the minimisation metric (see Methods).

**Figure SI 2:**
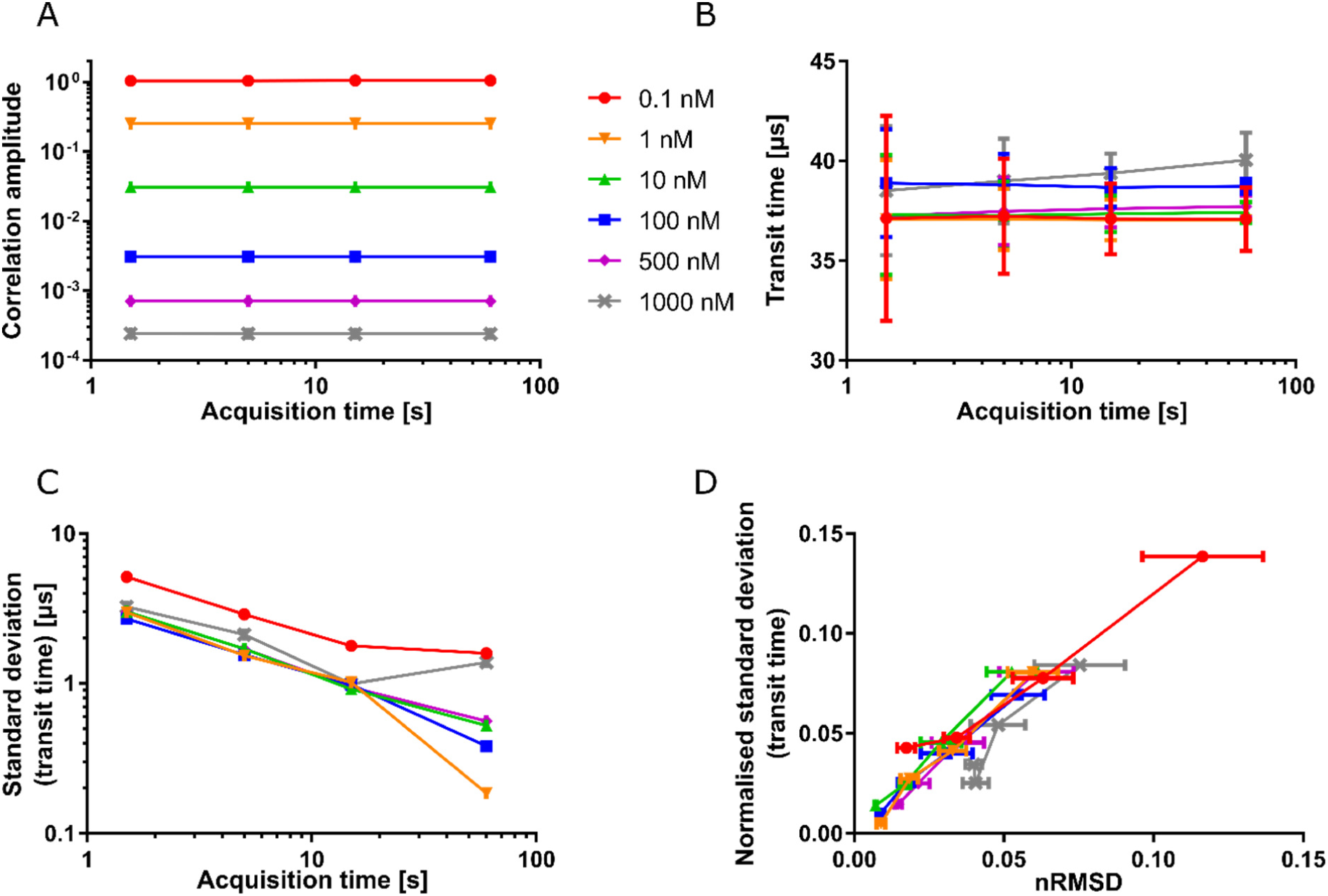
Effect of acquisition time on FCS performance at different concentrations of Atto655 in aqueous solution **A** The fitted correlation amplitude, **B** transit time, and **C** standard deviation of the fitted transit time, plotted against the acquisition time. **D** Relative standard deviation of the fitted transit times against the FCS noise level (nRMSD). Measurements were recorded at excitation laser power 15.4 µW and acquisition time 60 s, and then chopped into shorter intervals, as indicated.

**Figure SI 3:**
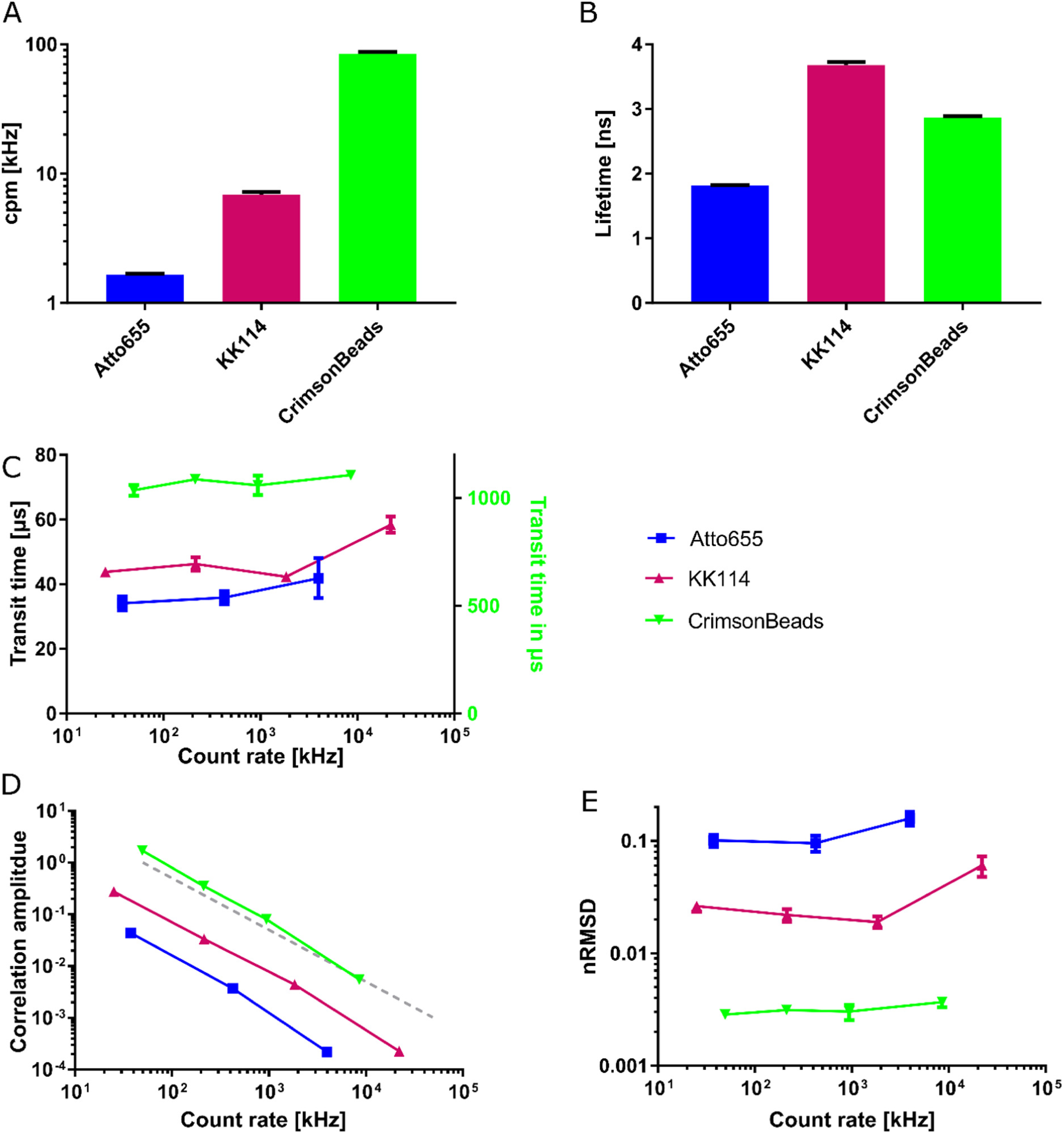
Influence of molecular brightness and fluorescence lifetime of dyes on performance in FCS at different count rates. **A** Counts per molecule and **B** fluorescence lifetime from single exponential fits to TCSPC histograms, for aqueous solutions of Atto655, KK114 (Abberior STAR Red) and Crimson Beads **C** Fitted transit time, **D** Correlation amplitude, and **E** FCS noise level (nRMSD) obtained for various concentrations of Atto655 (0.01, 0.1, and 1 µM), KK114 (0.01, 0.1, 1, and 10 µM) and Crimson Beads (1:2000, 1:500, 1:100, and 1:10 dilutions of the purchased stock solution), resulting in different count rates. Excitation laser power 1.5 µW, acquisition time 60 s, data-points represent averages and standard deviations of at least 3 measurements. The grey dashed line in **D** represents a linear relationship. Average nRMSD values from panel E were used for the inset in Figure 2E.

**Figure SI 4:**
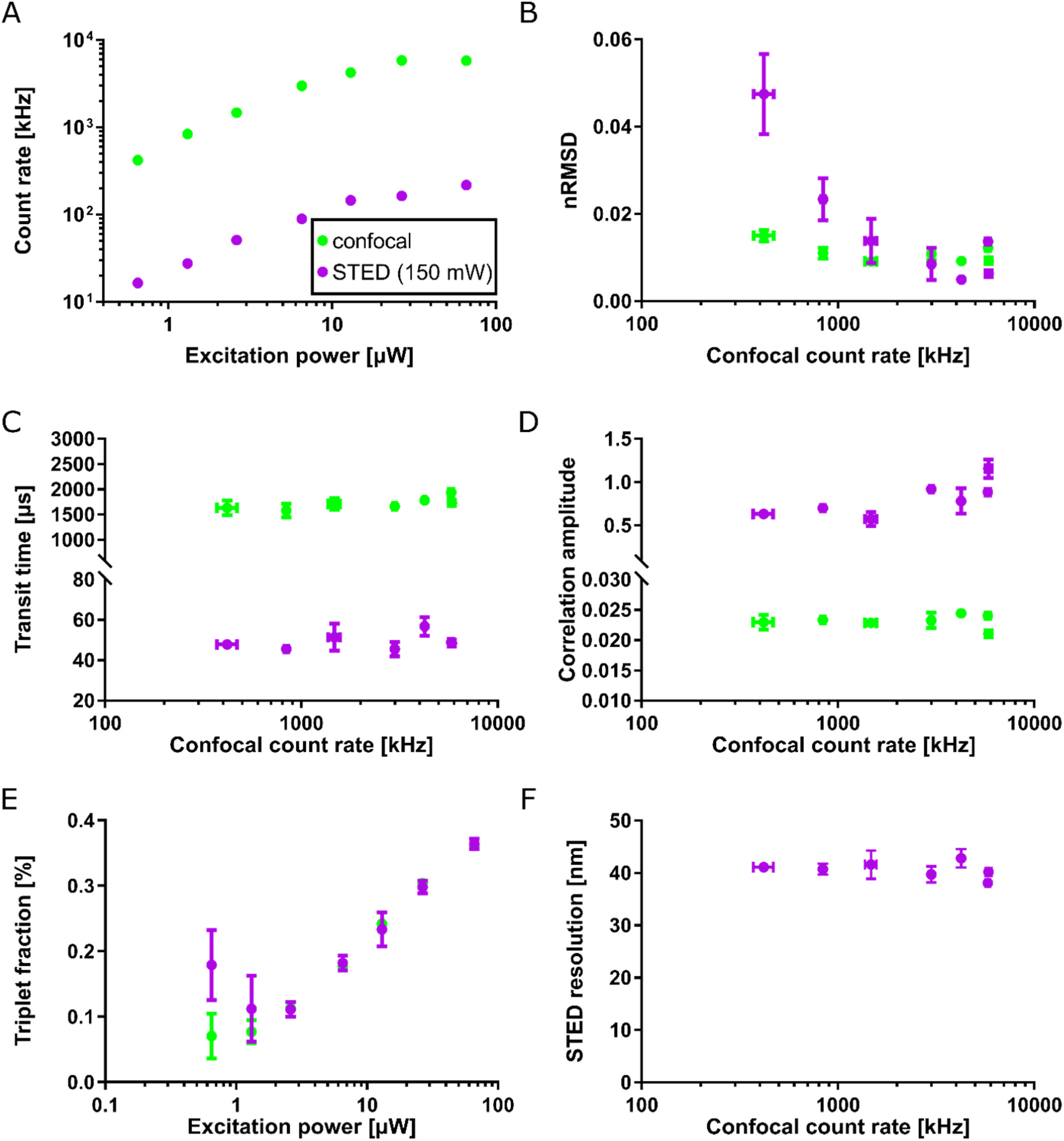
STED-FCS measurements of fluorescent Abberior STAR Red-DPPE in a DOPC SLB – acquired at varying excitation laser powers from 0.7 µW to 66 µW. **A** Change in confocal (green) and STED (purple) count rate when varying the excitation power. **B** The FCS noise level (nRMSD), and fit values of **C** transit times, **D** correlation amplitude, **E** triplet fractions, and **F** apparent STED resolution plotted against confocal count rate for confocal (green) and STED recordings (purple). Acquisition time 15 s, depletion laser power 150 mW. Data-points represent averages and standard deviations of at least 3 measurements.

**Figure SI 5:**
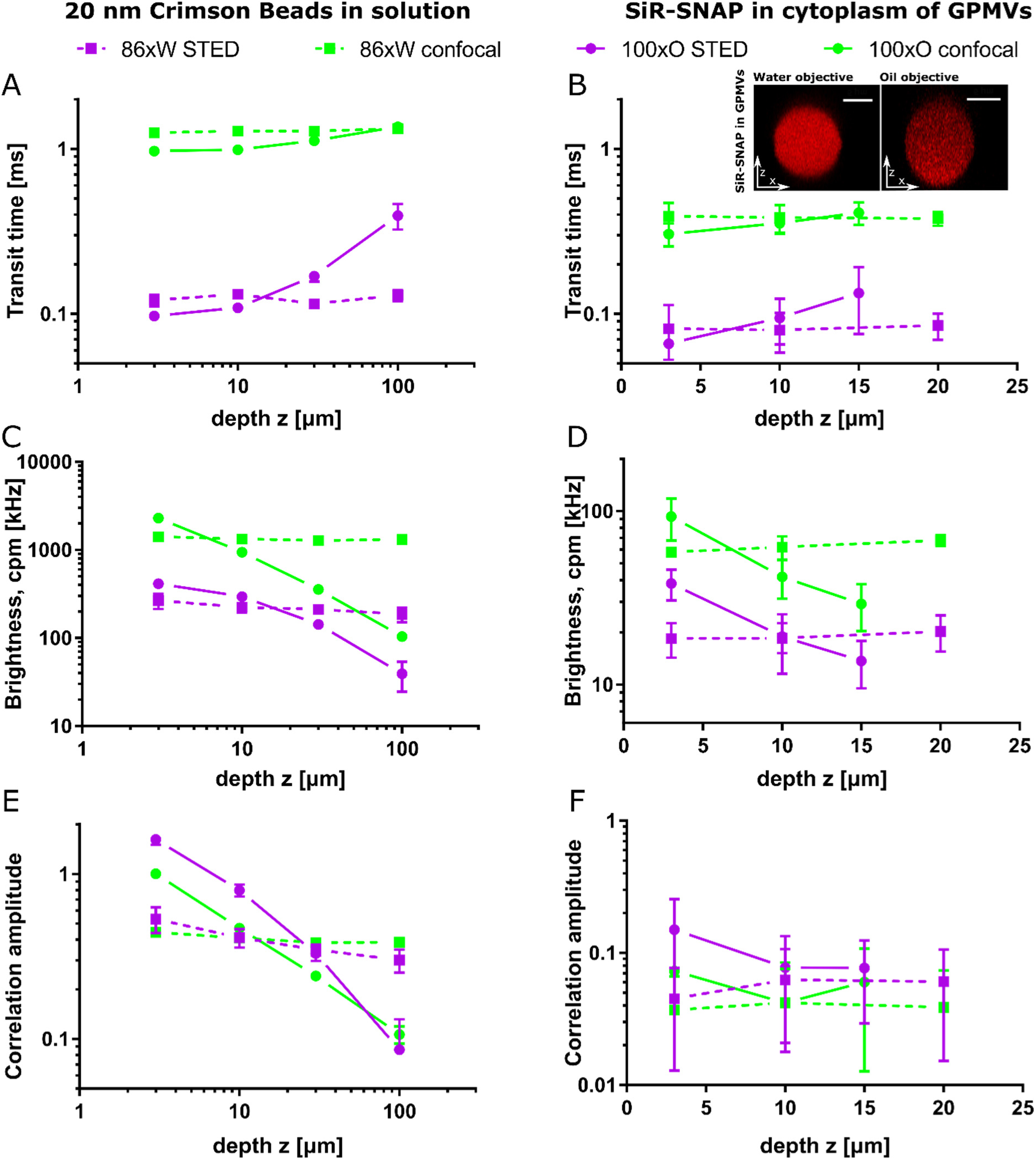
Performance of (STED-)FCS depends on objective and depth. (STED-)FCS experiments at different focal depth (z) for 20 nm Crimson beads in water **(left panel A, C, E)** and SiR-SNAP in cytoplasm of immobilised CHO GPMVs **(right panel B**,**D**,**F)** using a 100x oil- (circles, solid lines) and 86x water-immersion objective (squares, dashed lines). **A**,**B** The transit times, **C**,**D** molecular brightness (cpm) and **E**,**F** correlation amplitude determined from measurements at different focal depths in confocal (green) and STED mode (purple). Excitation laser power 8.9 µW, depletion laser powers 140 and 56 mW for crimson beads and SiR-SNAP, respectively, acquisition time 20 s. All data points are averages and standard deviations from at least three measurements. The inset in **B** shows confocal images of axial cross-sections of CHO GPMVs with SiR-SNAP in the cytoplasm.

**Figure SI 6:**
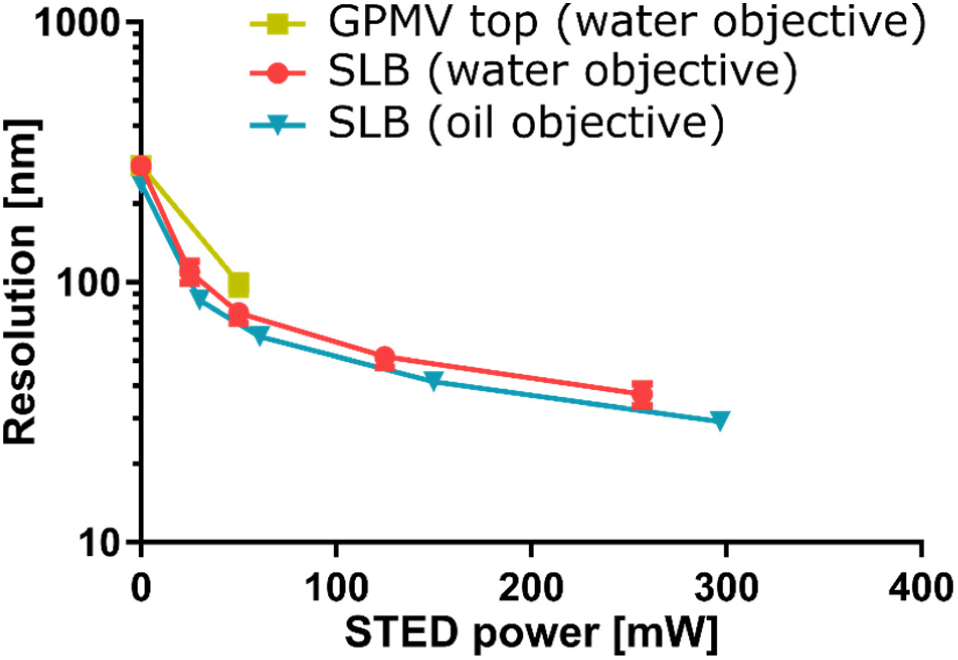
STED-FCS resolution calibration using SLBs (Abberior STAR Red-DPPE in DOPC) with oil- (blue) and water-immersion (orange) objectives, compared to the resolution obtained from the top membrane of CHO GPMVs (labelled with freely diffusing Abberior STAR Red-PEG-Cholesterol with a water-immersion objective (yellow)). Excitation power was 2.3 µW and depletion power was 125 mW, acquisition time 15 s. From at least three measurements per condition, the average value and their standard deviation are shown.

**Figure SI 7:**
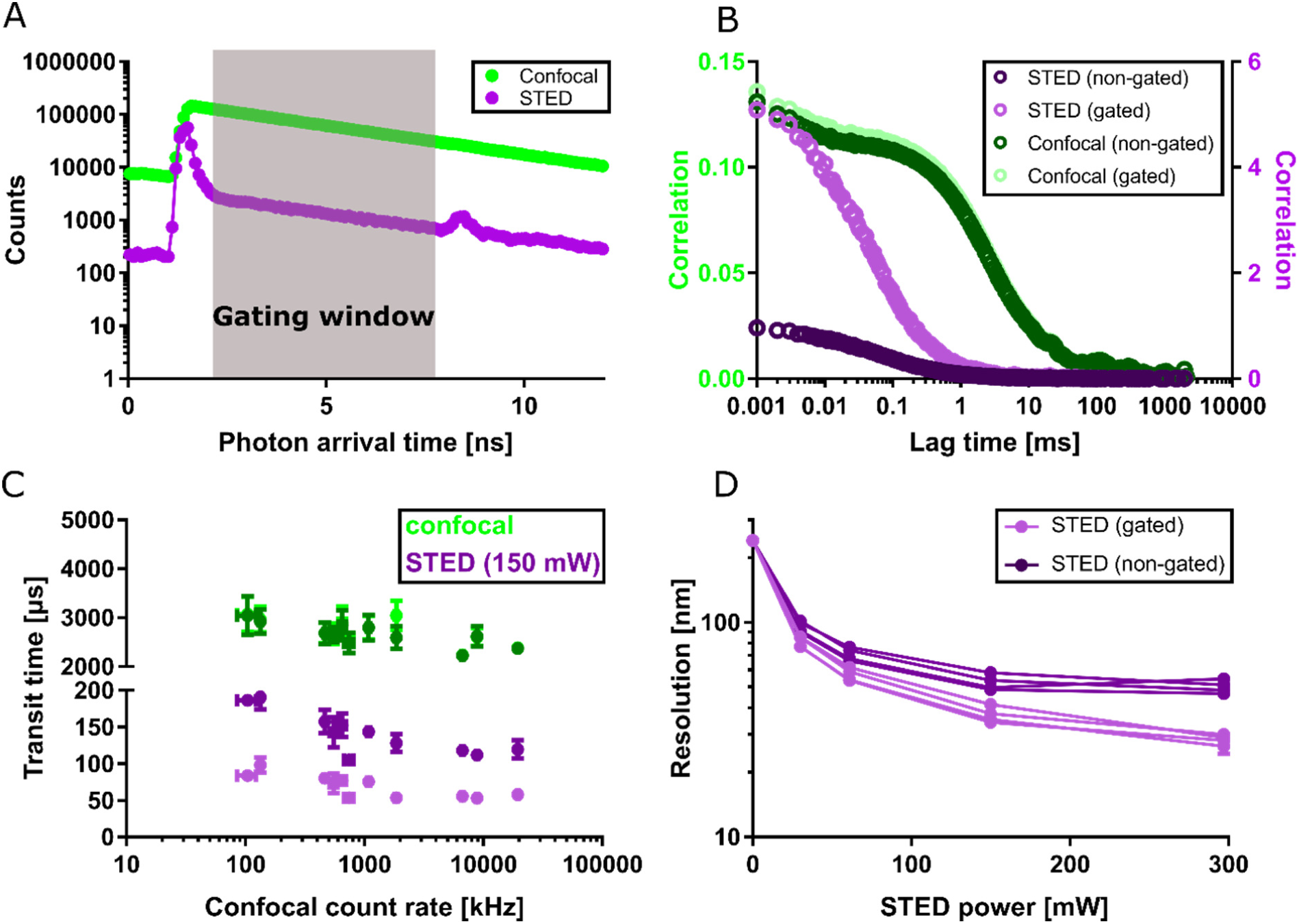
Influence of time gating in STED-FCS **A** Time correlated single photon counting (TCSPC) histograms of photon arrival times for an exemplary confocal (green) and STED-FCS measurement (purple, 150 mW depletion power) on a DOPC SLB labelled with Abberior STAR Red - DPPE. The gating window (grey area) was chosen to exclude confocal contributions to the STED-FCS correlation and to exclude the bump from the white light laser reflection in our system (note that it is present with the same absolute intensity in both datasets, but less visible for confocal data with higher fluorescence counts due to logarithmic scaling of the y-axis). **B** Exemplary correlation curves, and **C** average fitted transit times for confocal (green) and STED recordings (purple), analysed with and without gating applied (lighter and darker symbols, respectively). **D** Apparent resolution (i.e., diameter of the effective observation spot) as calculated from confocal and STED transit times for gated (light purple) and non-gated (dark purple) data acquired at different STED laser powers. Measurements of 15 s were acquired at 2.3 µW excitation laser power. In panels C and D, average and standard deviation of three measurements are shown.

**Figure SI 8:**
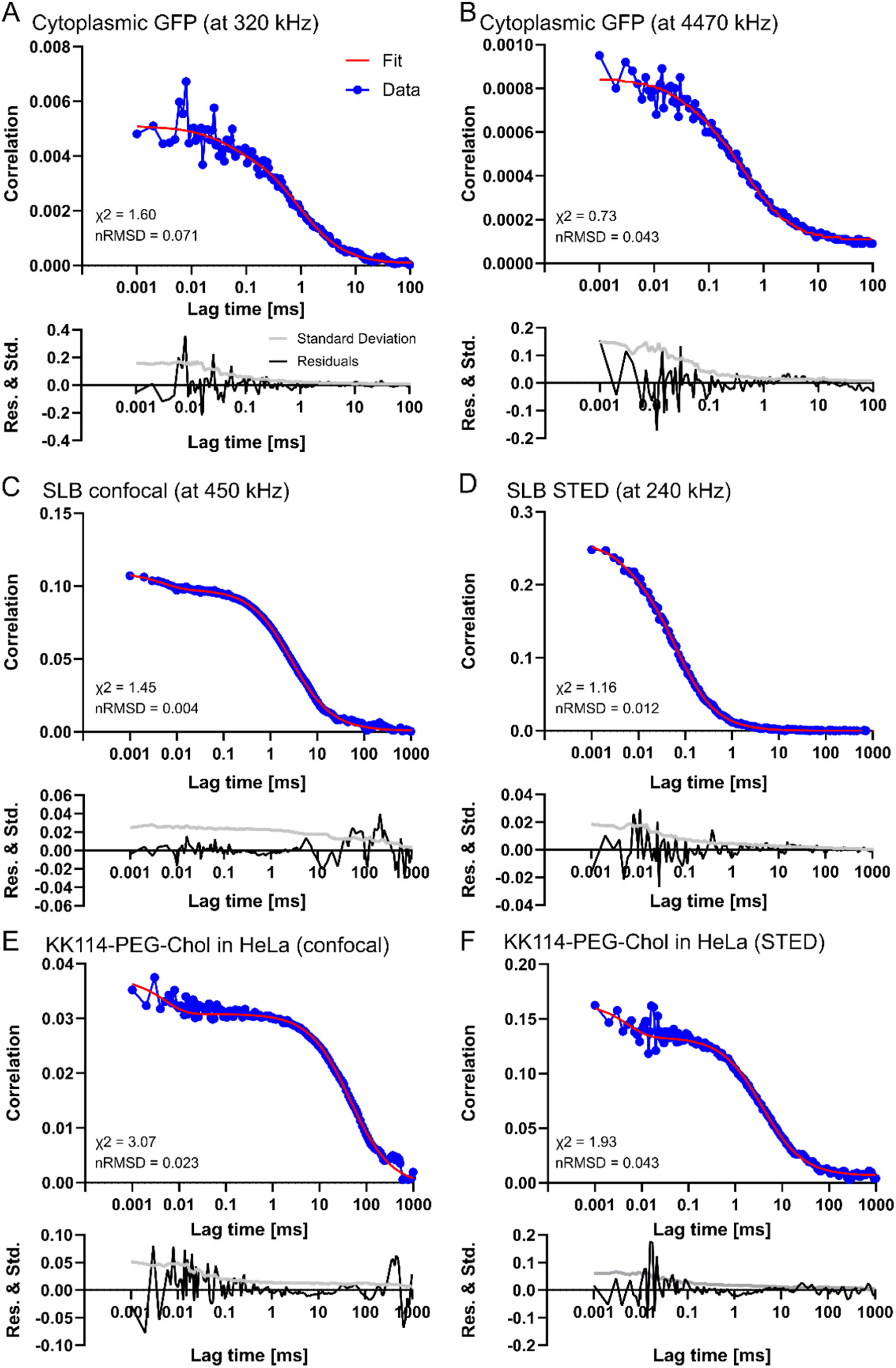
Representative FCS data with their fits (top panels), non-weighted fitting residuals (Res., black lines) and experimental standard deviation of the autocorrelation (Std., grey lines) plotted in the bottom panels, both normalised to the amplitude of the respective FCS curve. **A** and **B**: Fitting of a 3D diffusion model with a triplet state to correlation curves obtained from cytoplasmic GFP at high and low count rates, respectively (data from Figure 3). **C** and **D**: Fitting of a 2D diffusion model with a triplet state to confocal and STED data of KK114-DPPE diffusion in an SLB composed of DOPC (data from Figure 4). **E** and **F**: Fitting of a 2D diffusion model with a triplet state to correlation data obtained from KK114-PEG Cholesterol diffusing in the plasma membrane of live HeLa cells in confocal and STED acquisition mode (data from Figure 5). Non-weighted residuals were used to determine our signal quality measure nRMSD, whereas the standard deviations were taken into account to calculate the reduced χ^2^ as the minimisation metric in fitting (see Methods for details).

## Notes

#### Summary of Updates

Additional analysis, improved text

